# *FKBP5* expression is related to HPA flexibility and the capacity to cope with stressors in the house sparrow

**DOI:** 10.1101/2021.03.09.434659

**Authors:** Cedric Zimmer, Haley E. Hanson, Lynn B. Martin

## Abstract

The hypothalamic-pituitary-adrenal (HPA) axis and its end products, the glucocorticoids, are critical to responding appropriately to stressors. Subsequently, many studies have sought relationships between glucocorticoids and measures of health or fitness, but such relationships are at best highly context dependent. Recently, some endocrinologists have started to suggest that a focus on HPA flexibility, the ability of an individual to mount appropriate responses to different stressors, could be useful. Here, we tested the hypothesis that expression of *FKBP5*, a cochaperone in the glucocorticoid receptor complex, is a simple and reliable proxy of HPA flexibility in a wild songbird, the house sparrow (*Passer domesticus*). We quantified HPA flexibility in a novel way, using guidance from research on heart rhythm regulation. As predicted, we found that adult sparrows with low stress-induced *FKBP5* expression in the hypothalamus exhibited high HPA flexibility. Moreover, low *FKBP5* expression was associated with better stress coping capacities in terms of exploratory disposition and body mass maintenance. Altogether, these results suggest that *FKBP5* may be important in the regulation HPA flexibility, potentially affecting how individuals cope with natural and anthropogenic adversity.

## 1. Introduction

In nature, animals face numerous stressors, and appropriately responding to these dynamic and unpredictable challenges is crucial to fitness. Such responses usually require the ability to adjust the phenotype to match prevailing conditions (Taff & Vitousek, 2016; Zimmer, Hanson, Wildman, Uddin & Martin, 2020a). One system important for such adjustments is the hypothalamic-pituitary-adrenal (HPA) axis (Taff & Vitousek, 2016; Zimmer et al., 2020b). The HPA axis coordinates organismal responses to stressors mainly by regulating the secretion of glucocorticoids (primarily cortisol in most mammals and fish and corticosterone in birds, rodents, amphibians and reptiles) (Wingfield et al., 1998; Sapolsky, Romero & Munck, 2000). In the absence of stressors, circulating levels of glucocorticoids are maintained at low levels, mediating traits related to energy balance such as foraging activity, metabolism and body mass (Sapolsky et al., 2000). When facing stressors, however, rapid and dramatic increases in circulating glucocorticoids temporarily redirect energy away from non-essential activities and towards traits important to short-term survival (Wingfield et al., 1998; Sapolsky et al., 2000). These stress responses are critical to coping with unpredictability and adversity, as individuals that do not elevate glucocorticoids tend to be unable to adjust phenotypic responses appropriately (Darlington, Chew, Ha, Keil & Dallman, 1990; Thaker, Vanak, Lima & Hews, 2010). However, high glucocorticoid concentrations can be costly, resulting in physiological damage, accelerated senescence or even infectious disease (McEwen, 2008; Angelier, Costantini, Blévin & Chastel, 2018). Such costs do not only depend on maximal glucocorticoid concentrations but also on the duration of elevations and subsequent tissue exposure (Zimmer et al., 2019; Zimmer et al., 2020b). The duration of stress responses is thus tightly regulated by negative feedback mechanisms, which ultimately return glucocorticoids to baseline concentrations (Romero, 2004). Faulty negative feedback can be associated with low survival probability or reproductive success (Romero & Wikelski, 2010; Zimmer et al., 2019).

Although many empirical studies find relationships between glucocorticoid concentrations and measures of fitness, these relationships are at best highly context dependent and at worst non-existent (Schoenle, Zimmer & Vitousek, 2018; Schoenle, Zimmer, Miller & Vitousek, 2021). These inconsistencies may be due to the complexity of HPA axis regulation. As above, both increases and decreases in glucocorticoids comprise effective HPA regulation. Further, inconsistent results have led many researchers to begin to advocate for research on HPA flexibility itself, which we and others define as within-individual, rapid and reversible change in HPA regulation in response to unpredictable challenges (Taff & Vitousek, 2016; Zimmer et al., 2020a). To date, most researchers have used single or a few glucocorticoid measurements to try to capture HPA flexibility (Schoenle et al., 2018; Zimmer et al., 2020a). We question this approach, given the unlikelihood that a few hormone measures could describe such a complex trait. We argue that any measure of HPA flexibility must account for the most salient aspect of what the trait represents: a mechanism that enables individual organisms to realize optimal phenotypes quickly or adeptly via glucocorticoid regulation (Zimmer et al., 2020a; Zimmer et al., 2020b). Any proxy of HPA flexibility must thus entail repeated measurements of baseline, post-stressor, and negative feedback glucocorticoid concentrations in the same individuals.

When glucocorticoid concentrations are relatively high, as they are during stress response, their effects are mainly regulated through the binding of glucocorticoids to glucocorticoid receptors (GR). Whereas GR is expressed in nearly every cell type of the body (Cohen & Steger, 2017), it is the binding of glucocorticoids to GR in the major control sites of the HPA axis (i.e., the hypothalamus, hippocampus and anterior pituitary gland) that regulate the phenotypic response to elevated glucocorticoids. GR binding in these regions in particular leads to alterations of the expression of thousands of genes as well as negative feedback on HPA activity, collectively adjusting behavior and physiology to complement prevailing conditions (Landys, Ramenofsky & Wingfield, 2006; Dickens, Romero, Cyr, Dunn & Meddle, 2009; Cornelius et al., 2018). In this sense, any defensible form of HPA flexibility must comprise GR signaling (Zimmer et al., 2020a). In many contexts, however, circulating glucocorticoid concentrations do not reflect the extent of GR-mediated activation of glucocorticoid responsive elements within the genome (Haque, Mifsud, Price & Reul, 2021). This endocrinological disconnect casts yet more doubt on using simple or single circulating glucocorticoid measures as proxies of HPA flexibility. Perhaps no single, simple marker of such a complex trait as HPA flexibility exists, but one particular aspect has shown promise in human biomedical research, *FKBP5*, the gene encoding the FK506 binding protein 51 (Zimmer et al., 2020a).

FKBP5 is a cochaperone in the GR complex that has an inhibitory effect on GR signaling and activity. In humans and lab rodents, *FKBP5* expression is upregulated by elevated glucocorticoid concentrations, creating an ultrashort, intracellular negative feedback loop on GR activity that controls GR sensitivity to glucocorticoids. High *FKBP5* expression can also increase GR resistance resulting in an amplified stress response by reducing HPA axis negative feedback efficacy. Altogether, *FKBP5* expression controls GR activity and signaling, ultimately affecting the cellular and organismal response to stressors (Rein, 2016; Zannas, Wiechmann, Gassen & Binder, 2016). In lab mice, variation in *FKBP5* regulation was associated with individual differences in glucocorticoid stress responses and negative feedback efficacy (Touma et al., 2011; Hoeijmakers et al., 2014; Häusl et al., 2021). This variation was also related to anxiety-like behaviors and cognitive flexibility (Touma et al., 2011; Hoeijmakers et al., 2014; Sabbagh et al., 2014; Blair et al., 2019). For example, mice with low *FKBP5* expression showed an attenuated stress response and strong negative feedback associated with enhanced stress coping behavior (i.e., exploration) (Touma et al., 2011; Hoeijmakers et al., 2014). In light of HPA flexibility, therefore, *FKBP5* expression might capture an individual’s propensity for GR-mediated alterations of gene expression and by extension encode HPA flexibility (Lee et al., 2011; Menke et al., 2012; Rein, 2016; Zannas et al., 2016). We hypothesized that *FKBP5* expression is a fundamental element of HPA flexibility, effectively underpinning the ability of an individual to match its phenotype to prevailing stressors via glucocorticoids (Zimmer et al., 2020a). Such a role for *FKBP5* expression in humans and lab models is supported (Lee et al., 2011; Menke et al., 2012; Lee, 2016; Rein, 2016; Zannas et al., 2016), but research on *FKBP5* and HPA flexibility in wildlife is presently lacking. In the present study, we asked whether *FKBP5* is a viable proxy of HPA flexibility in a wild songbird, the house sparrow (*Passer domesticus*).

To determine whether *FKBP5* expression in the HPA axis is associated with HPA flexibility, we brought wild house sparrows into captivity and probed the effects of stressors on glucocorticoid regulation and *FKBP5* and GR expression in key regions of the HPA axis in the brain. Captivity alone can be a potent stressor for wild animals (Martin, Kidd, Liebl & Coon, 2011; Love, Lovern & DuRant, 2017; Fischer, Wright-Lichter & Romero, 2018). House sparrows lose weight for as long as nine weeks after introduction to captivity. Captivity also results in dysregulation of the immune system and the HPA axis, usually leading to high baseline glucocorticoid levels over extended periods (Martin et al., 2011; Love et al., 2017; Fischer et al., 2018). Decreases in body mass can result from diet shifts and modification of organs size, but it has also been attributed to increased glucocorticoids, as body mass loss is consistently observed in animals exposed to chronic stressors (Dickens & Romero, 2013; Love et al., 2017; Fischer et al., 2018). To maximize the amount of HPA flexibility we could observe among individuals, we exposed half of the birds in the study to a standard protocol consisting of three, 30-minute exposures to acute stressors (e.g., human disturbance, noise) daily for 20 days. This protocol alters HPA activity and other traits (e.g., behavior, body mass) in this and other songbird species (Rich & Romero, 2005; Cyr & Romero, 2007; Lattin & Romero, 2014; Gormally, Wright-Lichter, Reed & Romero, 2018). In all birds, we measured concentrations of the glucocorticoid corticosterone i) at baseline, ii) after 30 minutes of restraint, and iii) after treatment with dexamethasone (to induce negative feedback) four times over the course of several weeks. We then used these data to calculate HPA flexibility for each individual using an approach (RMSSD) initially described for heart rate variability (Figure 1). From most blood samples, we also measured *FKBP5* expression, and at the end of the experiment, we measured *FKBP5* and GR expression following a restraint protocol (stress-induced) in the hippocampus and hypothalamus, the two main regulatory regions of the HPA axis. Finally, we recorded changes in body mass over the course of the study as well as exploratory behavior and neophobia twice during the experiment as organismal endpoints that might be related to HPA flexibility and/or *FKBP5* expression. We predicted that individuals with lower stress-induced *FKBP5* expression in the HPA axis would show higher HPA flexibility and cope better with challenges (i.e., maintain body mass and be more exploratory) (Zimmer et al., 2020a). In previous studies, individual variation in HPA axis activity was associated with differences in exploratory disposition and neophobia (Cavigelli & McClintock, 2003; Zimmer, Boogert & Spencer, 2013). We also tested whether *FKBP5* expression in the blood was correlated with stress-induced expression in the hypothalamus and hippocampus, as it is in mice (Ewald et al., 2014; Lee, 2016), and whether GR and *FKBP5* were correlated in multiple tissues in the same individual birds.

**Figure 1:**
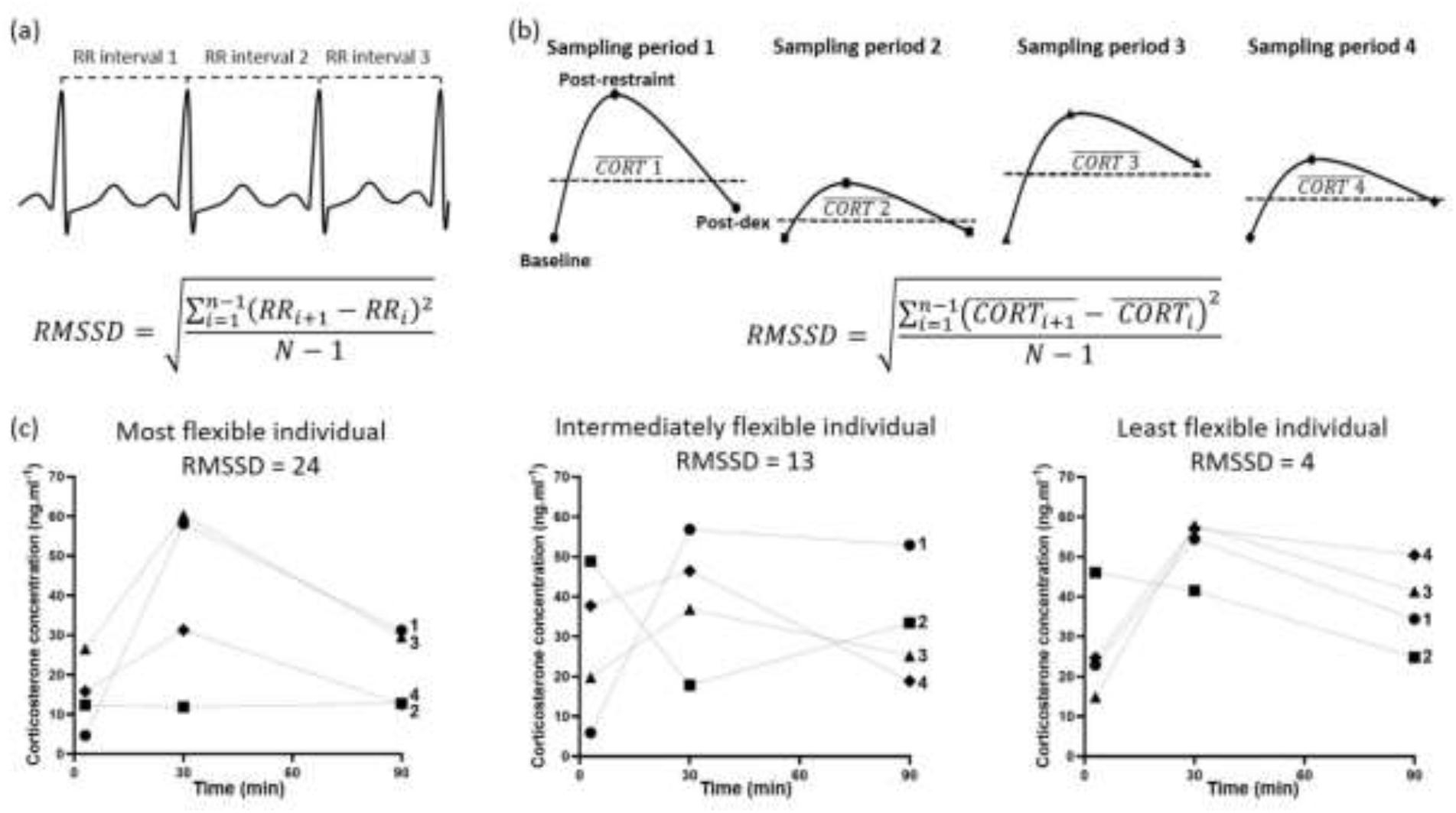
Calculation of HPA flexibility using the square root of the mean squared differences (RMSSD) of average corticosterone concentrations in house sparrows across several measurements of HPA activity. (a) Heart rate variability (HRV) analyses are based on variation in the intervals between successive heartbeats (RR interval). In this context, the most informative analysis is the RMSSD of successive RR intervals; higher RMSSD values indicate higher HRV. (b) We suggest that the RMSSD of successive measurements of mean corticosterone concentrations 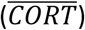 over successive sampling periods could be a meaningful measure of HPA flexibility. Here, we estimated RMSSD for all corticosterone concentrations over the entire study, calculating flexibility in average corticosterone concentrations across the three measurements in a sampling period (baseline, post-restraint, and post dexamethasone (post-dex)) over the four corticosterone sampling periods (1: capture (circle), 2: before the beginning of the chronic stress protocol (square), 3: halfway through the chronic stress protocol (triangle), 4: at the end of the chronic stress protocol (diamond)). Here, as with HRV, higher RMSSD values indicates higher HPA flexibility. (c) Representative individual corticosterone profiles over the four sampling periods: at capture (1, circle), before the beginning of the chronic stress protocol (2, square), halfway through the chronic stress protocol (3, triangle) and at the end of the chronic stress protocol (4, diamond). For illustrative purposes, we represent the bird with the highest RMSSD (24) that has the highest HPA flexibility, a bird with the median RMSSD value (14) that has intermediate HPA flexibility and the bird with lowest RMSSD (4) that has the lowest HPA flexibility.

## 2. Material and Methods

### 2.1 Sample collection and chronic stress protocol

We captured house sparrows in the Tampa Bay region, Florida, using mist nests in early July 2020. At capture in the field (sampling period 1), a blood sample was taken from each bird within 3 minutes of it contacting a net to measure baseline circulating corticosterone. A second blood sample was taken after each bird was held for 30 minutes in a cloth bag to measure post-restraint corticosterone. Immediately after this blood sample was taken, individuals were injected in the pectoral muscle with 1mg.kg^−1^ dexamethasone (60 μL of 0.5 mg.mL^−1^ dexamethasone sodium phosphate solution, Henry Schein) to assess HPA axis negative feedback efficacy (Gormally et al., 2018). A final blood sample was taken 60 minutes after dexamethasone injection to measure post-dexamethasone (post-dex) corticosterone concentration. Upon transfer to the University of South Florida vivarium, birds were randomly assigned to two groups: control (n=9, 2 adult females, 3 adult males, 4 juveniles) and chronic stress (n=9, 2 adult females, 2 adult males, 5 juveniles). Groups were housed in two different rooms (see supplementary material). Birds of both groups were exposed to the same sampling paradigm as in the wild thrice more (Figure 1b): i) the day prior to the chronic stress protocol (sampling period 2), ii) half-way through the chronic stress protocol (day 10, sampling period 3), and iii) the day after the final chronic stress protocol (sampling period 4). To ensure baseline concentrations were collected for all sparrows, a large group of researchers simultaneously entered the room in which birds were housed, successfully obtaining blood from all the birds within 4 minutes of entering the room. At each sampling period, individuals bleeding order was randomized to limit bias and variability in corticosterone profiles that could be introduced by the sampling regime.

All samples (about 50 μL) were collected in heparinized microcapillary tubes by pricking the brachial vein with a sterile needle. About 10 μL of blood was transferred into a microcentrifuge tube containing 300 μL of RNAlater (Ambion) and kept on ice until transferred at −80°C until RNA extraction. The rest of the blood was transferred into an empty microcentrifuge tube and kept on ice until centrifugation, then plasma was frozen at −20°C until hormone analysis. After the first blood sample for each of the four periods, the body mass of each bird was recorded (to 0.1g).

The chronic stress group was exposed to a 20-day standardized chronic stress protocol involving 30 minutes of acute stressors three times a day (except on day 10 when the middle stress series occurred) (Rich & Romero, 2005; Cyr & Romero, 2007; Lattin & Romero, 2014). The stressor type and time of administration were randomly determined, with at least 2 hours intervening each stressor. We used six different stressors that have been shown to alter corticosterone concentrations in this species and are intended to elicit mild to moderate psychological stress without physical discomfort (Rich & Romero, 2005; Lattin & Romero, 2014; Gormally et al., 2018) (see supplementary material for details). Control birds were not subjected to the above stressors and left otherwise undisturbed except for husbandry or for blood sampling. Following the last sampling period, all birds were euthanized (223 ± 25 min after initial disturbance) via isoflurane overdose and rapid decapitation. About 10 μL of blood was collected and stored in RNAlater. Brains were immediately removed using RNA-free tools, flash frozen on dry ice and stored at −80°C until microdissection.

### 2.2 Behavioral test

All birds were tested for neophobia and exploration of a novel environment before the start and just before the end of the chronic stress protocol. Tests occurred over two days with half of the birds of each group exposed to the neophobia test first and the other half to the exploration test first. Neophobia tests took place between 07:30 and 08:30. Food was removed from the cages 30 minutes before lights went off the night before the neophobia test, and birds tested early the following morning when they were motivated to approach a food dish. The latency to approach the food dish surrounded by a novel object (i.e., within a body length of distance from the dish) and to feed from the dish, and the number of approaches and feeds were recorded for each bird (see supplementary for detailed procedure).

Novel environment exploration tests took place between 08:30 and 11:30, alternating between birds from both groups. The novel environment was set up in a tent (Fabrill HQ200; 178 x 178 x 203 cm). Six remote controlled cabinet LED lights were glued to the tent ceiling to illuminate the tent. One-way mirror film was added to a window on the tent, allowing the experimenter to observe birds in the tent without being seen. The latency to first hop and/or flight as well as total hops and flights were recorded over a 10 minute period. The novel environment was also separated into four zones and the latency to enter each zone and the number of zones visited were quantified for each bird (see supplementary for detailed procedure).

### 2.3 Corticosterone assay

Corticosterone concentrations for all blood samples were determined using an enzyme immunoassay kit (DetectX Corticosterone, Arbor Assays: K014-H5) previously validated for house sparrows (see (Martin, Kilvitis, Thiam & Ardia, 2017) for details). The intra-assay variation based on duplicate samples was 4.89 % and the inter-assay variation based on a plasma pool run across plates was 7.15 %.

### 2.4 RNA extraction quantitative Real-Time PCR

We collected hippocampus and hypothalamus from each sparrow (see supplementary material for details). RNA was extracted from whole hippocampus and hypothalamus and 75 μL of blood/RNA later mixture for the first and final sampling periods using a TRI-reagent extraction method and was then diluted to 25 ng.μL^−1^ (Martin, Liebl & Kilvitis, 2015). From the resulting extracts, we measured *FKBP5* and GR (*NR3C1*) mRNA abundance in the hypothalamus and hippocampus, and *FKBP5* mRNA abundance in the blood using qRT-PCR (see supplementary for primers sequences and validation, and method details).

### 2.5 Statistical analyses

We used Generalized Linear Mixed Models (GLMM), fitted with a gamma distribution, to analyze the corticosterone data using proc GLIMMIX in SAS OnDemand (SAS Institute Inc.). We first ran a model on corticosterone data for sampling period 1 (capture from the wild samples) to determine whether HPA activity differed between adult and juvenile birds. Treatment group (stressors or controls), sex (female, male, juvenile), sample (baseline, post-restraint, post-dex) and their interactions were included as fixed factors. Individual identity was added as random factor. This analysis showed that HPA function differed between adults and juveniles but not between females and males (see results). Therefore, we included age (adult or juvenile) instead of sex as a factor for subsequent analyses. We then analyzed corticosterone changes across all four blood sampling periods using GLMM with treatment group, sampling period (blood samples at sampling period 1, 2, 3 and 4), bird age, sample (baseline, post-restraint, post-dex) and their interactions as fixed factors and individual identity as random factor. Post‐ hoc comparisons were performed using Tukey‐Kramer multiple comparison adjustments to obtain corrected p‐values.

We analyzed stress-induced *FKBP5* and GR relative expression in the hippocampus and hypothalamus using Generalized Linear Models (GLM) fitted with a gamma distribution using proc GENMOD. Each model included treatment group, age and their interaction as fixed factor. We used a GLMM fitted with a gamma distribution to analyze *FKBP5* relative expression in the blood with treatment group, age, sampling period (1 and 4), sample and their interactions as fixed factors and individual identity as a random factor. We estimated baseline *FKBP5* expression repeatability in adults across the four sampling period using a linear mixed model-based estimate of repeatability using the *rptr* package in R (4.0.3) (Stoffel, Nakagawa & Schielzeth, 2018). We added sex and treatment as control variables. Confidence intervals were estimated with parametric bootstrapping (1,000 iterations). Data were log transformed to reach normality.

We finally investigated potential relationships among *FKBP5* relative expression across tissues and GR relative expression, HPA flexibility and stress coping capacities using proc CORR to perform Spearman correlations. For relationships that appear to be driven by statistical outliers, we calculated Cooks D. We probed correlations separately in adults and juveniles. As there is no existing quantitative measure of HPA flexibility, we developed one based on a method used to describe heart rate variability (HRV; Figure 1). HRV analyses are mainly focused on variation in the duration between successive heartbeats, and a popular and informative descriptor of this variability is captured by the square root of the mean squared differences (RMSSD) of subsequent heartbeat intervals. RMSSD for HRV is typically calculated by first quantifying the time difference between subsequent heartbeats, then squaring differences among many successive heartbeats and averaging differences before obtaining the square root of the total. RMSSD thus measures flexible adjustments of heart rates to context (e.g., breathing, vagal tone, stress, etc.) (Figure 1a) (Park et al., 2017; Shaffer & Ginsberg, 2017). Here, we adapted RMSSD for corticosterone concentrations, assuming that the RMSSD of the mean corticosterone concentration of each sampling period over the four sampling periods would capture meaningful interindividual variation in HPA flexibility. Figure 1b describes our approach in detail and reveals the suitability of RMSSD as a proxy of HPA flexibility (Figure 1c).

As measures of stress coping capacities, we used body mass change between the first and final measurements and indexes of exploratory disposition and neophobia for each bird (Zimmer et al., 2013). For the behavioral data, we used proc PRINCOMP to perform principal component analyses (PCA) based on correlation matrixes of all individual behaviors for each behavioral assay after removing collinear variables, providing us with one exploratory and one neophobia index for each bird. For exploration behavior, we ran a PCA including the latency to move (hop or fly), the number of moves (hops and flights) and the number of zones visited. The first principal component (PC) accounted for 71% of the variation with an eigenvalue of 2.12 and PC2 explained 21% of the variance with an eigenvalue of 0.62. For neophobia, we ran a PCA including the latency to approach and feed and the number of approach and feed. PC1 explained 57% of the variance with an eigenvalue of 2.29 and PC2 explained 25% of the variance with an eigenvalue of 0.99. Therefore, we used each PC1 score as our exploration and neophobia indexes in later Spearman correlations with *FKBP5* expression. For significant correlations, we performed a model selection to determine if the significant *FKBP5* expression measure was a better predictor of behavioral performance than corticosterone concentrations measures (see supplementary).

## 3. Results

### 3.1 Corticosterone concentrations

HPA activity can differ between adults and juveniles and also change rapidly as birds mature (Wada, 2008; Bebus, Jones & Anderson, 2020; Jones, Nguyen & DuVal, 2020), so we first checked whether corticosterone concentrations at capture (sampling period 1) differed among females, males and juveniles. As expected, corticosterone concentrations differed among females, males and juveniles during this first sampling period (F_4,36_ = 3.99, p = 0.003, Figure S3). Baseline and post-restraint corticosterone concentrations were lower in juveniles than adults (t ≥ 2.46, p ≤ 0.018, Figure S3) but did not differ between adult females and males (t ≤ 1.57, p ≥ 0.15, Figure S3). Post-dex corticosterone concentrations in adult males were higher than juveniles (t = 2.16, p = 0.036, Figure S3) whereas adult females did not differ from either group (t ≤ 1.41, p ≥ 0.18, Figure S3).

As HPA activity at capture differed between adults and juveniles but not between females and males, we analyzed all subsequent data with the factor, age (adult *vs*. juveniles). Corticosterone concentrations for the different samples differed between treatment groups and series (group x series x sample: F_9,199.6_= 3.65, p = 0.0003, Figure S4, S5). Multiple comparisons did not show specific differences between the control and chronic stress groups. However, in both groups, baseline corticosterone was lower at capture (1: control 11.65 ± 7.75 ng.ml^−1^; stress 10.70 ± 6.42 ng.ml^−1^) than in the three other sampling periods (2: control 51.34 ± 18.91 ng.ml^−1^; stress 23.85 ± 15.78 ng.ml^−1^; 3: control 16.05 ± 10.37 ng.ml^−1^; stress 23.48 ± 13.66 ng.ml^−1^; 4: control 21.80 ± 10.57 ng.ml^−1^; stress 17.44 ± 10.77 ng.ml^−1^; t ≥ 1.97, p ≤ 0.049, Figure S4, S5).

### 3.2 Effects of age and stressor treatment on gene expression

In the hippocampus, stress-induced *FKBP5* and GR expression were not affected by stressor treatment (*FKBP5*: control 0.81 ± 0.40, stress 0.93 ± 0.33; GR: control 0.82 ± 0.30, stress 0.87 ± 0.19; χ_1,18_ ≤ 1.64, p ≥ 0.20, Figure S6) or age (*FKBP5*: adult 0.96 ± 0.41, juvenile 0.77 ± 0.33; GR: adult 0.82 ± 0.28, juvenile 0.86 ± 0.22; χ_1,18_ ≤ 2.83, p ≥ 0.09, Figure S6). In the hypothalamus, though, *FKBP5* expression was influenced by both stressor treatment and age (treatment x age: χ_1,18_ = 22.37, p < 0.0001, Figure S6) with juveniles in the chronic stress group (2.37 ± 0.52) expressing more *FKBP5* than control juveniles (0.92 ± 0.11) and more than adults in both groups (control 1.21 ± 0.12; stress 1.20 ± 0.46) (t ≥ 4.73, p ≤ 0.0001, Figure S6). However, GR expression did not differ between treatment (control 0.96 ± 0.29, stress 1.16 ± 0.29; χ_1,18_ = 2.88, p = 0.09) or age (adult 1.02 ± 0.28, juvenile 1.10 ± 0.33; χ_1,18_ = 0.11, p = 0.74) groups.

In the blood, *FKBP5* expression in the different samples differed across sampling periods (1 and 4) (sample x series: F_2,108_ = 4.66, p = 0.012, Figure S7). Baseline and post-restraint *FKBP5* expression did not differ between the first (baseline: 0.0018 ± 0.0009, post-restraint: 0.0016 ± 0.0010) and the last (baseline: 0.0022 ± 0.0014; post-restraint: 0.0023 ± 0.0020) sampling period. Within each sampling period baseline and post-restraint expression did not differ (t ≤ 1.23, p ≥ 0.87, Figure S7). Within each period, *FKBP5* expression in the post-dex sample (1: 0.0040 ± 0.0017; 4: 0.0096 ± 0.0074) was higher than baseline and post-restraint samples (t ≥ 5.11, p < 0.0001, Figure S7). Expression in the post-dex sample was higher in the final sampling period than at capture (t = 4.68, p = 0.0002, Figure S7). When repeatability was examined in adults, *FKBP5* expression in baseline blood samples across the four sampling periods was repeatable (R = 0.45 ± 0.18 [0.04-0.71], p = 0.003).

### 3.3 Relationships between *FKBP5* expression and HPA flexibility

In adults (Table S1), hypothalamic stress-induced *FKBP5* expression was negatively correlated with RMSSD (−0.77, p = 0.016); individuals with low *FKBP5* expression had high HPA flexibility (Figure 2). Similarly, *FKBP5* expression in baseline blood samples taken at initial capture was also negatively correlated with HPA flexibility (−0.85, p = 0.004, Figure 3a). *FKBP5* expression in the hippocampus and other blood samples were not related to HPA flexibility (Table S1). In juveniles (Table S2), we did not observe any relationships between *FKBP5* expression and HPA flexibility.

**Figure 2:**
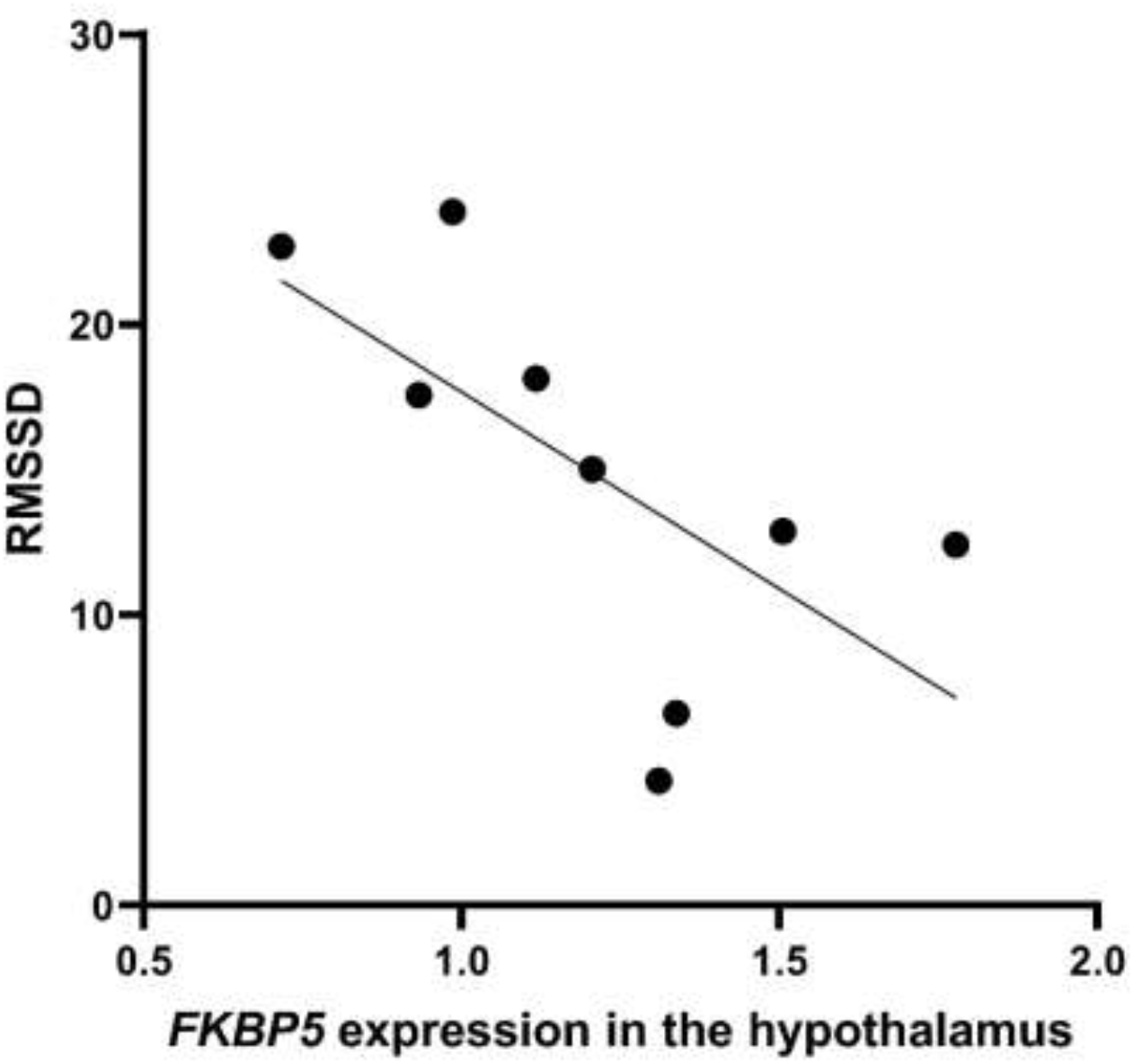
Relationships between *FKBP5* expression and HPA flexibility in house sparrows. Relationships between the square root of the mean squared differences (RMSSD) of successive corticosterone sampling period and *FKBP5* expression in the hypothalamus in adult sparrows.

**Figure 3:**
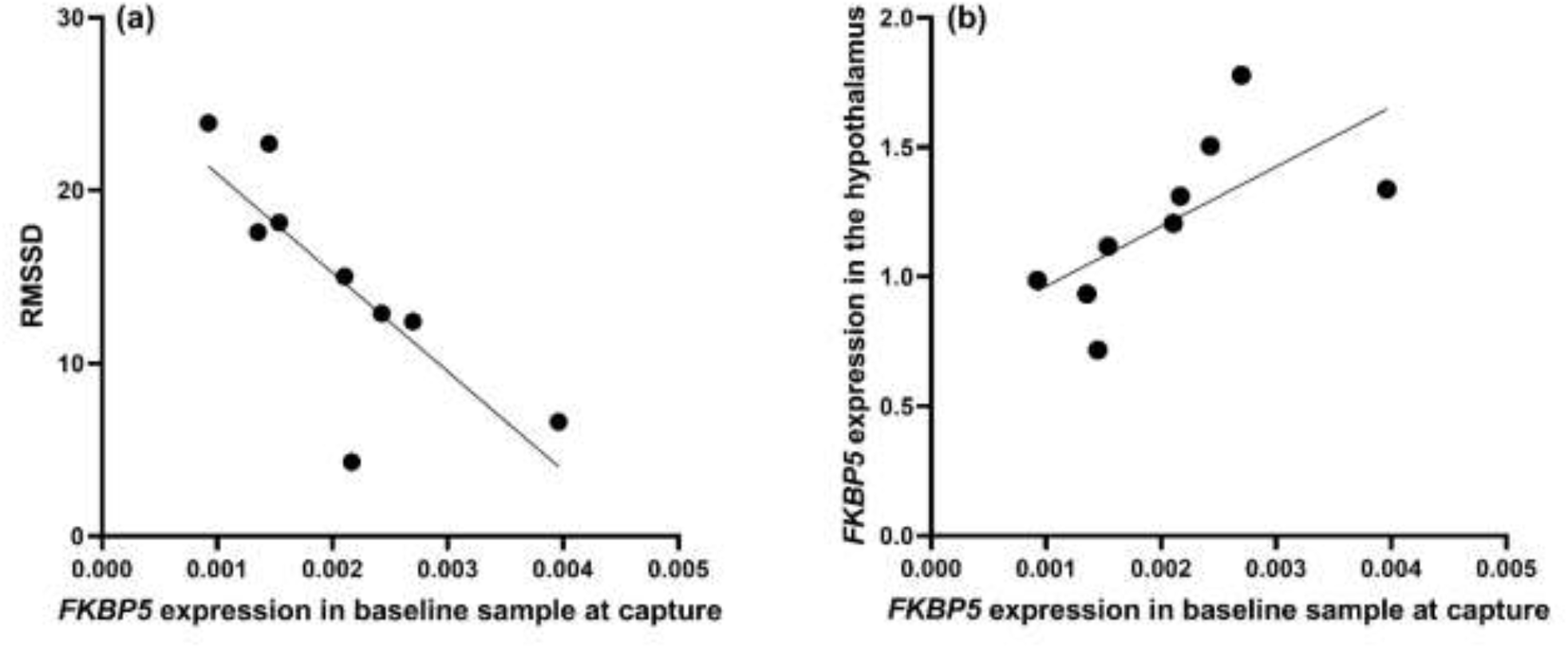
*FKBP5* expression as a potential biomarker of HPA flexibility. (a) Relationships between the square root of the mean squared differences of successive stress series (RMSSD) and *FKBP5* relative expression in baseline blood sample at capture in adults. (b) Relationship between stress-induced *FKBP5* expression in the hypothalamus and in the baseline blood sample at capture in adults.

### 3.4 Relationships between *FKBP5* expression and stress coping capacities

In adult sparrows (Table S1), hippocampal stress-induced *FKBP5* expression was negatively correlated with the exploration index at the first test (r = −0.67, p = 0.05); birds with lower expression were more exploratory of the novel environment (Figure S8a). *FKBP5* expression in the hypothalamus and blood samples was not related to any stress coping measures (Table S1). In juveniles (Table S2), *FKBP5* expression in the post-dex sample at capture was negatively correlated with neophobia index at the first test (r = −0.71, p = 0.033); juveniles with low *FKBP5* expression were less neophobic (Figure S8b). *FKBP5* in the post-dex sample of the final sampling period was negatively correlated with body mass change over the course of the study (r = −0.67, p = 0.05); juveniles with low *FKBP5* expression lost less body mass (Figure S8c). *FKBP5* expression in the hypothalamus, hippocampus and other blood samples was not related to any stress coping measures (Table S2). For the correlations between *FKBP5* expression with neophobia and with body mass loss in juveniles, a Cooks D test revealed that one statistical outlier influenced the relationships. To explore further the relationships between *FKBP5* expression and behavior and determine if *FKBP5* expression is a better predictor of performance in these tests than corticosterone concentrations, we used model selection, fitting a set of models and identifying best-fit models using AICc. For both behaviors, the model that included *FKBP5* expression had more support than any model including a measure of corticosterone concentration (exploration in adults: model weight including *FKBP5* expression in the hippocampus: 0.86, ΔAICc from the second model: 4.6 (Table S3), β = −2.2 ± 0.5; neophobia in juveniles: model weight including *FKBP5* expression in the post-dex sample at capture: 0.81, ΔAICc from the second model: 5.8 (Table S4), β = −190.1 ± 62.7).

### 3.5 Gene expression correlations among tissues and samples

In adults (Table S5), *FKBP5* expression was not correlated between hippocampus and hypothalamus (r = −0.13, p = 0.73), but within each tissue, *FKBP5* and GR expression were correlated (hippocampus r = 0.65, p = 0.05; hypothalamus r = 0.65, p = 0.05; Figure S9a). *FKBP5* expression in the hypothalamus was also correlated with *FKBP5* expression in baseline blood samples at initial capture (r = 0.88, p = 0.002; Figure 3b). In juveniles (Table S6), hippocampal *FKBP5* expression was correlated with hypothalamic expression (r = 0.82, p = 0.007; Figure S9b). *FKBP5* and GR expression were correlated in the hypothalamus (r = 0.83, p = 0.005; Figure 9c) but not in the hippocampus in juveniles. *FKBP5* expression was not correlated between brain and blood samples in juveniles.

## 4. Discussion

As hypothesized, stress-induced *FKBP5* expression in the hypothalamus was related to HPA flexibility (RMSSD) in adult house sparrows such that individuals with low *FKBP5* expression showed high flexibility. In juveniles, stress-induced *FKBP5* expression in the hypothalamus and hippocampus was unrelated to HPA flexibility, perhaps because of the immaturity of the HPA axis. In both adults and juveniles, our results suggest that low *FKBP5* expression is associated with better stress coping capacities, in terms of body mass maintenance, exploratory disposition, and neophobia. Overall, low *FKBP5* expression was associated with high HPA flexibility, which may result in greater stress coping capacity in these sparrows.

### 4.1 HPA flexibility and *FKBP5* expression in adult house sparrows

Based on existing biomedical research, we predicted a relationship between HPA flexibility and *FKBP5* expression in the hypothalamus, although no work to our knowledge in that field or wildlife endocrinology has ever explicitly demonstrated links between *FKBP5* expression and any measure of HPA flexibility (Zimmer et al., 2020a). Whereas RMSSD as a proxy of HPA flexibility requires more study, our results reveal expected relationships including with *FKBP5* expression in the blood at the time of capture. Here, adult sparrows with low stress-induced hypothalamic *FKBP5* expression had high HPA flexibility. We expected this relationship because *FKBP5* upregulation by exposure to stressors decreases GR affinity for glucocorticoids, increasing GR resistance and consequently reducing downstream GR-dependent gene expression (Lee, 2016; Rein, 2016; Zannas et al., 2016). In other words, as HPA flexibility inherently requires GR activity, we expect that *FKBP5* is probably related to HPA flexibility as it regulates how glucocorticoids and GR interact to produce phenotypic change. In that sense, *FKBP5* expression might implicate the semiotic information content (the functional information encoded in a signal) of circulating glucocorticoids that underpin HPA flexibility (Zimmer et al., 2020a). In support here, adult sparrows with low stress-induced *FKBP5* expression in the hippocampus, a critical region for situation appraisal, behavioral adaptation to stress during novel situations, and activity and cognition regulation (de Kloet, Vreugdenhil, Oitzl & Joëls, 1998; Gray, Kogan, Marrocco & McEwen, 2017), were more exploratory in a novel environment. We should note, though, that our sample size was modest and behavior and gene expression were measured about three weeks apart. However, as stress-induced *FKBP5* expression in the hippocampus better predicted exploration than any glucocorticoid concentration, our interpretation is reasonable. More work is of course necessary to understand *FKBP5* as a determinant of the semiotic information content of glucocorticoids, and an especially promising study design to do so would involve the study of animal behavior and *FKBP5* expression across a shorter time scale.

### 4.2 Age-dependency of FKBP5 relationships with HPA flexibility

In juvenile birds, stress-induced *FKBP5* expression in the hypothalamus and hippocampus was not related to HPA flexibility. This outcome is probably partly due to the immaturity of the HPA axis in this group. Indeed, HPA activity is comparatively blunted in young, altricial species. During early development, the HPA axis of nestlings is hyporesponsive to stressors, but as birds develop, baseline and post-restraint corticosterone concentrations increase to adult values (Figure 2) (Wada, 2008; Bebus et al., 2020; Jones et al., 2020). *FKBP5* expression is sensitive to early life conditions in rodents; exposure to early-life adversity permanently elevated *FKBP5* expression, which were related to traits used as diagnostics of mental health disorders for humans (Zannas et al., 2016). Here, juvenile birds exposed to the chronic stress protocol showed increased *FKBP5* expression in the hypothalamus, which could disrupt its effects on HPA flexibility. Additionally, as the juvenile HPA axis in passerines is relatively undeveloped, HPA flexibility might not yet be canalized in many birds, explaining the absence of a relationship with *FKBP5* expression among tissues. *FKBP5* expression is quite labile, modifiable by DNA methylation (Zannas et al., 2016; Wiechmann et al., 2019), particularly during early development (Zannas et al., 2016). It would be interesting in the future to determine how exposure to adversity during early-life affects DNA methylation and expression of *FKBP5* and their consequences for HPA flexibility in natural contexts. In lab model organisms, *FKBP5* expression is upregulated after exposure to a challenge, increasing GR resistance and decreasing GR signaling (Touma et al., 2011; Hoeijmakers et al., 2014; Rein, 2016; Zannas et al., 2016). *FKBP5* expression was somewhat labile in blood samples with an expected increased expression 90 minutes post-challenges (Menke et al., 2012; Wiechmann et al., 2019), which may explain why the post-dex expression is correlated with stress coping capacities. Early-life experience could thus work through *FKBP5* to affect stress coping capacity (via HPA flexibility). Perhaps modest increases in *FKBP5* expression in response to stressors might prime stress coping capacities of some genotypes (Zimmer et al., 2020a). Of course, the effects of an outlier in correlations for juveniles and the comparatively small number of adult and juvenile birds necessitate more work.

### 4.3 HPA flexibility measure and effects of stressors in captivity

Despite increasing appeals in the literature to study HPA flexibility (Taff & Vitousek, 2016; Zimmer et al., 2020a), we do not know of any other quantitative measure of HPA flexibility than the one we offer here. Our method was derived from a common measure of flexible adjustments of heart rates to context (Park et al., 2017; Shaffer & Ginsberg, 2017), and seemingly, it captures important variation in HPA flexibility among individuals (Figure 1). We therefore advocate for RMSSD as a measure of HPA flexibility in other systems. As birds in our study were exposed to different contexts between sampling periods, we expected that individuals with high HPA flexibility would exhibit different corticosterone profiles between successive sampling periods and conversely, individuals with low HPA flexibility would show more similar successive corticosterone profiles. Our data indeed show that birds with higher HPA flexibility have high RMSSD values whereas birds with low HPA flexibility have low RMSSD values (Figure 1c). Nevertheless, the use of RMSSD as a measure of HPA flexibility needs to be studied in more detail. Particularly, we used mean corticosterone for each measurement period to calculate HPA flexibility, but other elements such as the shape of glucocorticoid response profile (Figure 1), could be used instead.

One particularly important topic warranting study is whether high HPA flexibility enables quick matching of organismal phenotypes to current conditions (Taff & Vitousek, 2016; Zimmer et al., 2020a). Here, while lower *FKBP5* expression was related to higher HPA flexibility and the few behaviors we evaluated in captivity, we cannot conclude that high HPA flexibility is adaptive. Nevertheless, *FKBP5* expression in pre-captivity baseline blood samples was correlated with stress-induced expression in the hypothalamus and with HPA flexibility (Figure 3). The same type of correlation was found in lab mice after four weeks of corticosterone treatment (Ewald et al., 2014). However, in mice the blood was collected at the same time than the brain at the end of the experiment. Here, the relationship existed between expression in the hypothalamus at the end of the experiment following a restraint protocol and in pre-captivty baseline blood samples collected almost a month earlier. Considering the strength of the relationship despite our limited sample size (r= 0.88) and cellular heterogeneity between these tissues, it seems unlikely that this result is an artifact. While this relationship is somewhat surprising because of the time lag between the measures and the potential effect of captivity and treatment (chronic stress and dexamethasone), *FKBP5* expression in baseline blood samples across the sampling periods was appreciably repeatable, and baseline *FKBP5* expression at capture and at the end of the experiment were correlated. However, as final baseline expression was not correlated with expression in the hypothalamus, future work to rule out low sample size limiting statistical power are necessary. While our data suggest that *FKBP5* expression in the blood has the potential to be a marker of HPA flexibility in natural populations, it will be necessary to investigate further if this relationship exists using more samples in more species, ideally in natural contexts. Unfortunately, it is not possible to obtain baseline *FKBP5* expression in tissues of the HPA axis and HPA flexibility in the same individual, which prevents the direct study of the relationship between them. As *FKBP5* expression is labile, determining whether baseline or stress-induced *FKBP5* expression in the HPA axis are correlated with expression in the blood of individuals in natural setting would allow for more confidence in using baseline *FKBP5* expression in the blood as a marker of baseline and/or stress-induced *FKBP5* expression in HPA axis tissues.

Captivity can be a potent stressor for some wild species, altering body mass, immune functions and HPA axis regulation for weeks to months (Martin et al., 2011; Love et al., 2017; Fischer et al., 2018). To increase the likelihood we would reveal variation in HPA flexibility in the sparrows we studied, we exposed half the birds here to a chronic stress protocol. This protocol altered HPA axis regulation and stress coping capacity in other studies (Cyr & Romero, 2007; Lattin & Romero, 2014; Gormally et al., 2018), but here we found no effects of stressor exposure on corticosterone concentration or *GR* or *FKBP5* expression in the hippocampus and blood. We cannot rule out that this lack of effect may be due to the close proximity in time between sampling periods and/or the repeated use of dexamethasone. Only *FKBP5* expression in the hypothalamus was affected by treatment and age, but juveniles exposed to the chronic stress protocol showed higher *FKBP5* expression than adults and control juveniles. These results indicate that the stressor protocol had some but likely a modest effect, but also suggest that exposure to adverse conditions during HPA axis maturation may have long term consequences for *FKBP5* regulation and expression in wildlife.

## 5. Conclusion

We found that HPA flexibility was related to *FKBP5* expression in the hypothalamus of adult house sparrows. This relationship did not exist in juvenile birds probably because their HPA axes were immature. In both adult and juvenile sparrows, lower *FKBP5* expression also seems to have been associated with better stress coping capacity. Overall, these results suggest that *FKBP5* expression may play a crucial role in regulating HPA flexibility affecting how individuals cope with adverse conditions. In order to increase the likelihood to see variation we brought individuals in captivity and exposed some to chronic stress, which prevent to make any conclusion about the adaptive potential of high HPA flexibility. As a next step, it will be important to investigate whether these relationships hold in natural conditions and in particular, that lower *FKBP5* expression allows better coping with natural stressors. These studies will require measurements of *FKBP5* expression in blood and tissues of the HPA axis in non-stressed, wild individuals. Ultimately, it would be critical to test if *FKBP5* expression and regulation permit to adapt to rapid changing conditions and are associated with fitness.

## Acknowledgments

We thank Meredith Kernbach and Emily Ruhs for their assistance for capture and sampling. The authors thank members of the Martin lab at the University of South Florida for thoughtful feedback.

