## Supplementary material for "*FKBP5* expression is related to HPA flexibility and the capacity to cope with stressors in the house sparrow"

### Methods

#### *Capture and housing*

All birds were captured between 06:50 and 07:30 then transferred to the University of South Florida vivarium where they were individually housed in 35.6 x 40.6 x 44.5 cm cages for 28 days within visual and auditory contact of each other. Birds were housed under 14L:10D light conditions to match natural day length. Water and food (mixed seeds) were provided *ad libitum* at all-times except when food was removed just before neophobia tests.

#### *Chronic stress protocol*

We used six different stressors (See supplementary material for details). restraint in an opaque bag for 30 minutes; cage rolling where the cages were placed on lab carts and gently rolled back and forth for 30 min; cage disturbance where one person entered the housing room every 3 minutes and tapped on or rattled the front, top or sides of the cages for 30 seconds; exposure to a radio tuned to a local music station at a modest volume for 30 minutes; crowding where 5/4 birds were placed into one cage for 30 minutes; and human voice/presence in the room for 30 minutes. No stressor was used twice in the same day.

#### *Behavioral test*

Neophobia tests took place between 07:30 and 08:30. Food was removed from the cages 30 minutes before lights went off the night before the neophobia test, and birds tested early the following morning when they were motivated to approach a food dish. At the beginning of each test, the food dish was put back in the cages with a novel object made of a colorful bristle block surrounding the sides of the food dish or placed on the side of the dish (Figure S1). We used different objects of different colors between the two tests. Tests started as soon as the food dish was back in the last cage and birds were observed for 20 minutes by CZ from a corner of the room.

Novel environment exploration tests took place between 08:30 and 11:30, alternating between birds from both groups. The novel environment was set up in a tent (Fabrill HQ200; 178 x 178 x 203 cm). Six remote controlled cabinet LED lights were glued to the tent ceiling to illuminate the tent. One-way mirror film was added to a window on the tent, allowing the experimenter (CZ) to observe birds in the tent without being seen. Plastic panels were taped to the floor of the tent, creating separations, and 5 perches, 3 platforms and various novel objects made of bristle blocks were scattered around the tent floor. The position of the panels, perches and platforms, and object types differed between the initial and final tests (Figure S2). Each individual to be tested was caught from its cage and immediately transported to the testing room. Each bird was introduced alone to the novel environment with the room and tent lights off. Then, the experimenter took his position and turned on the light in the tent. Each bird had 5 minutes to acclimate to the tent before the experimenter started recording individual behavior for 10 minutes.

*RNA extraction and qTR-PCR*

Brains were placed on an upside-down, RNAse free petri dish on ice to thaw and then were dissected using anatomical landmarks, following previously established methods in dark-eyed juncos (*Junco hyemalis*) (Rosvall et al. 2012), to collect the hippocampus and hypothalamus. Brain samples were placed in microtubes on dry ice as soon as they were dissected and kept at -80°C until RNA extraction.

Primers for *FKBP5* (forward primer: TTTGAGAAGGCCAAGGAGTCGT; reverse primer: AGCCATACTCCATTTCCAGCCA) and GR (forward primer: TGAAGAGCCAGTCCCTGTTCGAG; reverse primer: CAACCACATCATGCATAGAGTCCAGCA (Banerjee et al. 2011). We previously validated *HMBS* as an appropriate housekeeping gene in this sparrow population (Hanson et al. submitted) and checked that its expression was not affected by stressor treatment. *FKBP5* and *GR* primers were validated using serial dilution of hypothalamus and hippocampus samples and RNA melt curves were checked for to ensure a single product was amplified. Melt curves revealed amplification of intended targets, a 103% efficiency for *FKBP5* and 105% efficiency for *GR*, with no apparent dimer formation.

All qRT-PCR reactions (20 µL) were run in duplicate alongside non-template controls (NTC) and no reverse transcriptase controls (NRT), on a Rotor-Gene Q system (Qiagen). Each well contained 10 µL of iTaq Universal SYBR Green One-Step Kit (Bio-Rad), 0.6 µL of forward primer, 0.6 µL of reverse primer, 0.25 µL of SCRIPT, 6.55 µL of nuclease free water and 2 µL of diluted RNA or 2 µL of nuclease free water for NTCs. For NRTs, reverse transcriptase was replaced by nuclease free water. Reactions entailed the following conditions: 10 minutes at 50°C for reverse transcription reaction, then 1 minute at 95°C for polymerase activation and DNA denaturation, followed by 40 amplification cycles of 15 seconds at 95°C then 30 seconds at 60°C. Melt-curve analyses were performed from 65 to 95°C with 0.5°C increment step every 3 seconds. A calibrator, a mix of RNA from 3 hippocampus and 3 hypothalamus samples of control birds, was run on all plates to calculate mRNA abundance using the comparative Ct method ($2^{-\Delta\Delta Ct}$), which reports mRNA abundance for each gene of interest as the fold change in expression compared to a calibrator sample and normalized to an internal reference gene (Livak and Schmittgen 2001).

Statistical analyses

We asked what was the best predictor of performances in the novel environment in adults and in neophobia in juveniles between the *FKBP5* expression measure that was correlated with performances in these test and measures of corticosterone concentration at capture. We chose to use corticosterone concentrations at capture because these measured occurred 3 or 4 days before the behavioral tests and were not affected by captivity. For both GLM model sets, corticosterone measures used were baseline, post-restraint and post-dex corticosterone concentrations as well as stress response (post-restraint – baseline) and negative feedback (post-dex – post-restraint). Best-fit models were identified by comparing the corrected Akaike information criterion (AICc) scores of the candidate models. For exploration in adults, all models included treatment, sex and their interaction as fixed effects. For neophobia in juveniles, all models included treatment as fixed effect.

**Supplementary figure and tables**

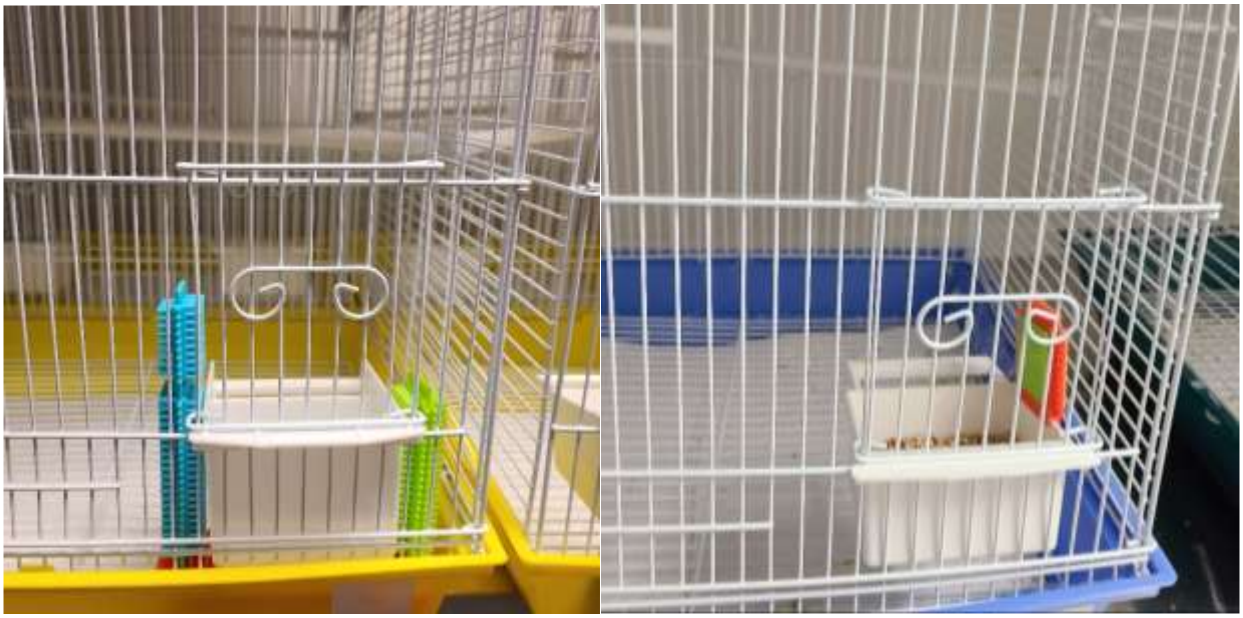

Figure S1: Photos of the objects used for the neophobia test before the start of the chronic stress protocol (left) and at just before the end of the chronic stress protocol (right).

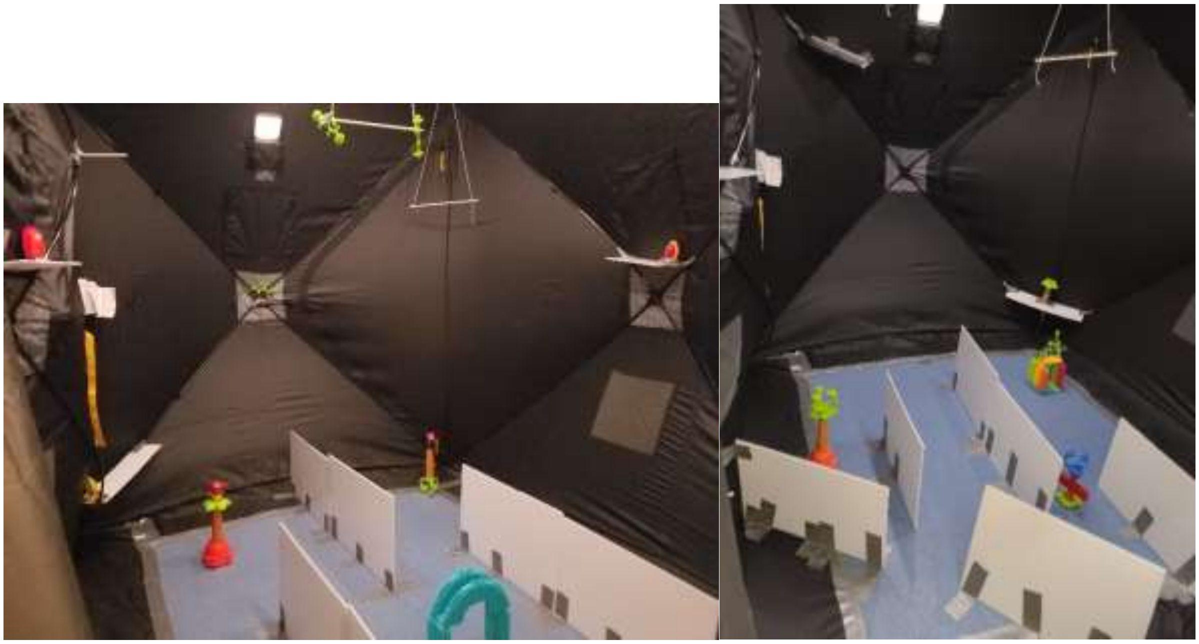

Figure S2: Photo of the inside of the tent used for the novel environment exploration test before the start of the chronic stress protocol (left) and at just before the end of the chronic stress protocol (right).

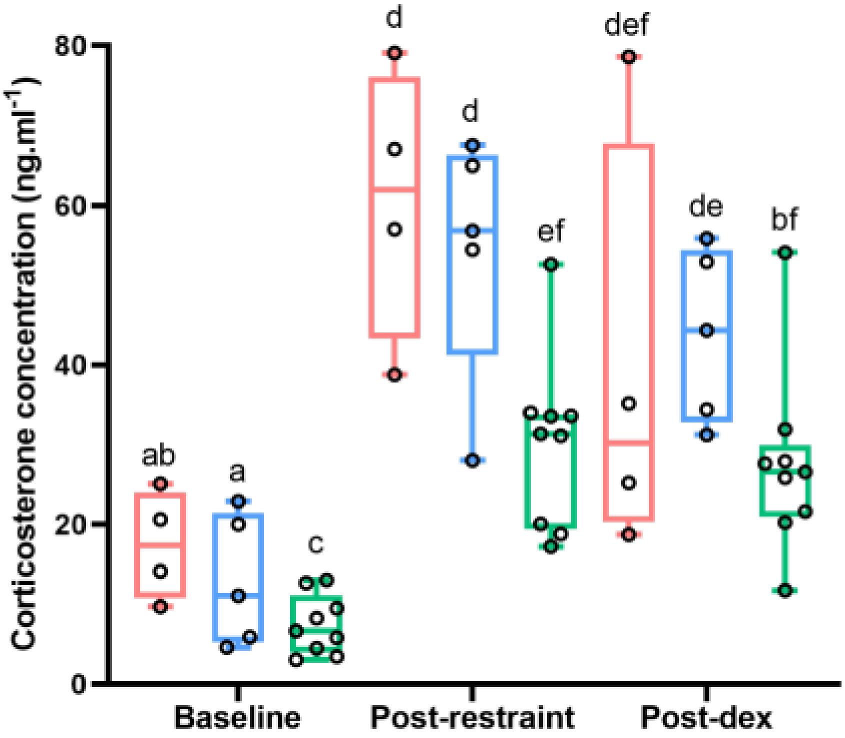


Figure S3: Corticosterone concentrations at capture**.** Baseline, post-restraint and post-dex corticosterone concentrations in females (red), males (blue) and juvenile sparrows (green) at capture. Different letters indicate significant pair-wise differences.

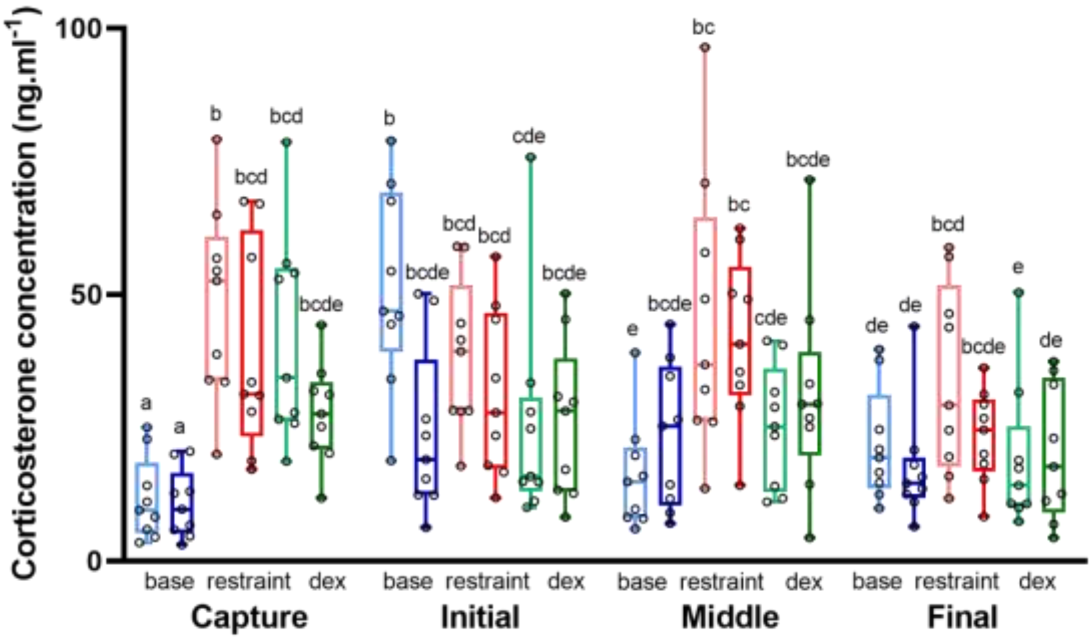


Figure S4: Corticosterone concentration for the four series. Baseline (base, blue), stress-induced (stress, red) and post-dex (dex, green) corticosterone concentration for the control group (light color) and the chronic stress group (dark color) for the four stress series. Different letters indicate significant differences.

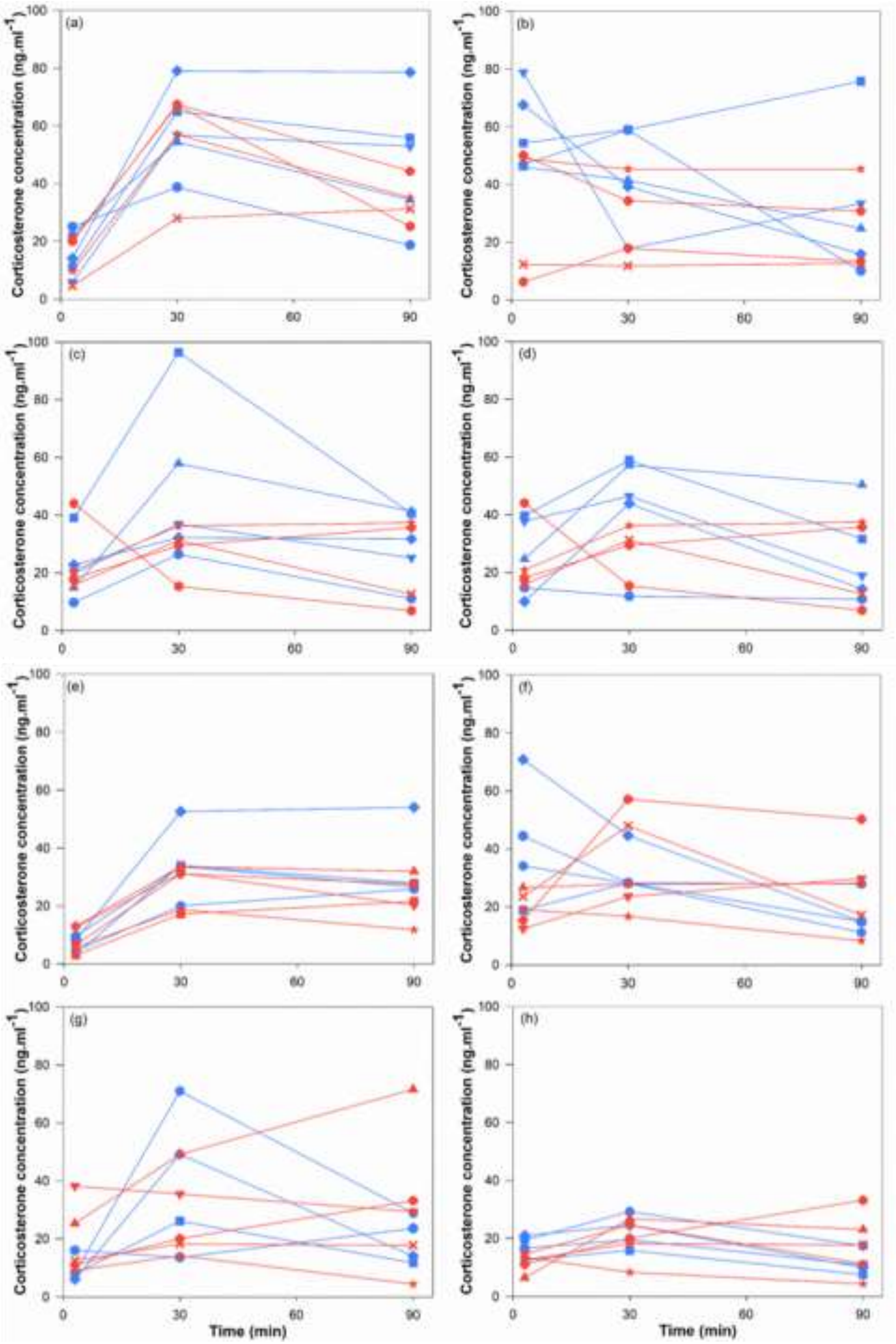


Figure S5: Individual corticosterone profiles at capture in adults (a) and juveniles (e), before the beginning of the chronic stress protocol (adults: b, juveniles: f), half way through the chronic stress protocol (adults: c, juveniles: g) and at the end of the chronic stress protocol (adults: d, juveniles: h) for the control group (blue) and chronic stress group (red). A different shape represents individuals for each age category.

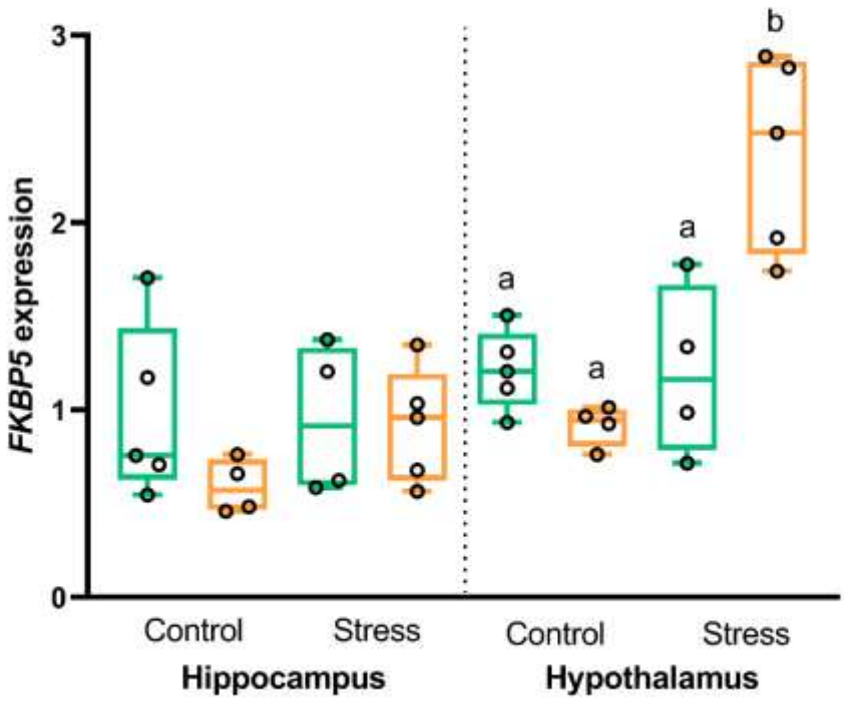


Figure S6: *FKBP5* relative expression in the brain. *FKBP5* relative expression in the hippocampus and hypothalamus of adults (green) and juveniles (orange) in the control and chronic stress groups. Different letters indicate significant differences between groups within the hypothalamus. *FKBP5* relative expression was significantly higher in hypothalamus than hippocampus.

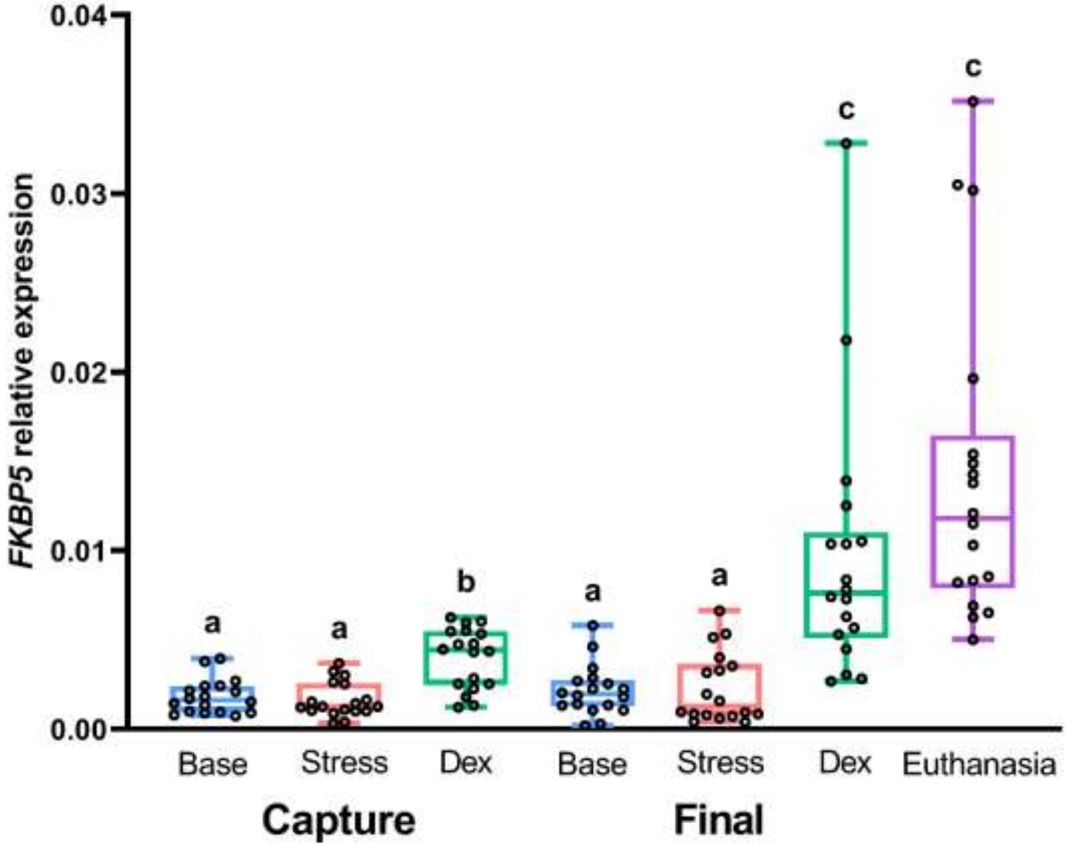


Figure S7: *FKBP5* relative expression in blood. *FKBP5* relative expression in blood at baseline (Base, blue), stress-induced (Stress, red) and post-dex (Dex, green) time point at capture and final series, and just before euthanasia (purple). Different letters indicate significant differences.

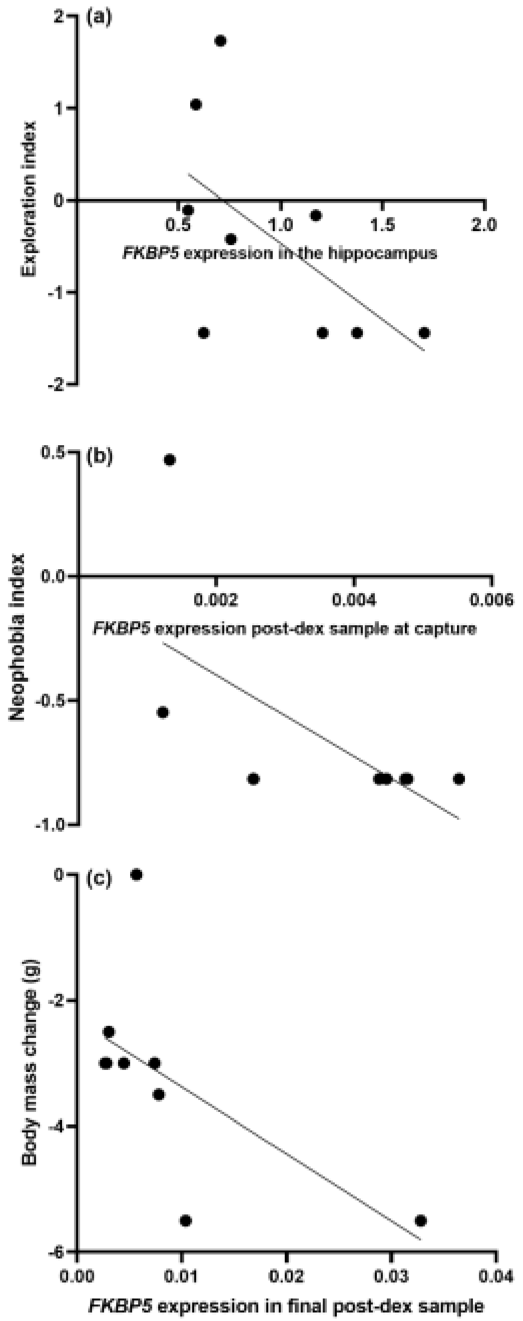


Figure S8: Relationships between *FKBP5* expression and measures of stress coping capacity. (a) Relationship between hippocampal *FKBP5* expression and exploration behavior index at the first test (higher in value indicates higher exploration) in adult sparrows. (b) Relationship between *FKBP5* expression in the post-dex blood sample at capture and neophobia at the first test (higher values indicate lower neophobia) in juveniles sparrows. (c) Relationship between *FKBP5* expression in the final post-dex sample and body mass change between the end of the experiment and capture in juveniles.

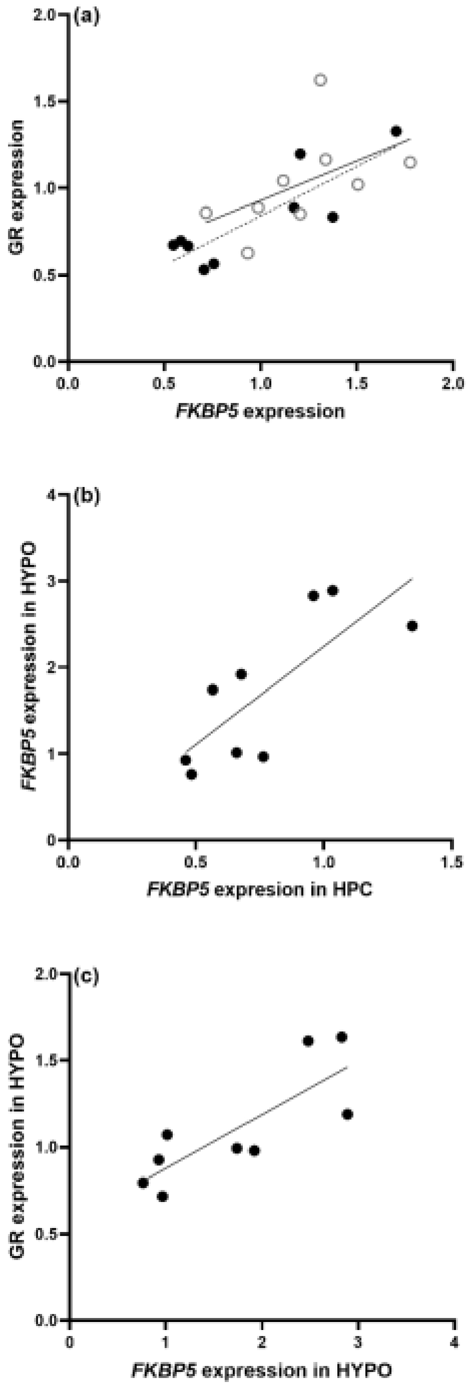


Figure S9: *FKBP5* expression relationships between tissues and with GR expression**.** (a) Relationships between *FKBP5* and GR expression in the hippocampus (closed circle, dash line) and hypothalamus (open circle, full line) in adults. (b) Relationships between *FKBP5* expression the hippocampus and hypothalamus in juveniles. (c) Relationship between *FKBP5* and GR expression the hypothalamus in juveniles.

Table S1: *FKBP5* relative expression in the brain (hippocampus (HPC) and hypothalamus (HYPO), in the blood baseline (base), stress-induced (str) and post-dex (dex) samples at capture (capt) and final series (final) relative expression in the brain correlation with index of HPA axis flexibility, body mass change between the end of the experiment and capture and index of neophobia and exploration in adults.

| | ***FKBP5* HPC** | ***FKBP5* HYPO** | ***FKBP5* base capt** | ***FKBP5* str capt** | ***FKBP5* dex capt** | ***FKBP5* base final** | ***FKBP5* str final** | ***FKBP5* dex final** |
|---|---|---|---|---|---|---|---|---|
| **RMSSD** | -0.45000<br>0.2242 | -0.76667<br>**0.0159** | -0.85000<br>**0.0037** | -0.43333<br>0.2440 | -0.31667<br>0.4064 | -0.20000<br>0.6059 | 0.08333<br>0.8312 | -0.21667<br>0.5755 |
| **Body mass change final - capture** | 0.40510<br>0.2794 | 0.24475<br>0.5256 | 0.43041<br>0.2475 | 0.00844<br>0.9828 | 0.42197<br>0.2579 | 0.12659<br>0.7455 | 0.47261<br>0.1989 | 0.27850<br>0.4680 |
| **Neophobia index initial** | 0.20084<br>0.6044 | 0.27615<br>0.4720 | 0.37657<br>0.3178 | -0.04184<br>0.9149 | -0.06695<br>0.8641 | 0.15900<br>0.6828 | 0.26778<br>0.4860 | 0.65273<br>0.0567 |
| **Neophobia index final** | 0.05941<br>0.8793 | 0.21783<br>0.5734 | 0.42576<br>0.2532 | 0.02970<br>0.9395 | -0.04951<br>0.8994 | 0.16833<br>0.6651 | -0.39606<br>0.2913 | 0.09901<br>0.7999 |
| **Exploration index initial** | -0.66462<br>**0.0506** | 0.46131<br>0.2113 | 0.25241<br>0.5123 | 0.08704<br>0.8238 | -0.13056<br>0.7378 | -0.16537<br>0.6707 | 0.07833<br>0.8412 | 0.18278<br>0.6379 |
| **Exploration index final** | -0.21667<br>0.5755 | 0.55000<br>0.1250 | 0.25000<br>0.5165 | 0.23333<br>0.5457 | 0.03333<br>0.9322 | 0.05000<br>0.8984 | -0.38333<br>0.3085 | -0.21667<br>0.5755 |

Table S2: *FKBP5* relative expression in the brain (hippocampus (HPC) and hypothalamus (HYPO), in the blood baseline (base), stress-induced (str) and post-dex (dex) samples at capture (capt) and final series (final) relative expression in the brain correlation with index of HPA axis flexibility, body mass change between the end of the experiment and capture and index of neophobia and exploration in juveniles.

| | ***FKBP5*** **HPC** | ***FKBP5*** **HYPO** | ***FKBP5*** **base capt** | ***FKBP5*** **str capt** | ***FKBP5*** **dex capt** | ***FKBP5*** **base final** | ***FKBP5*** **str final** | ***FKBP5*** **dex final** |
|---|---|---|---|---|---|---|---|---|
| **RMSSD** | 0.63333<br>0.0671 | 0.48333<br>0.1875 | 0.11667<br>0.7650 | 0.55000<br>0.1250 | -0.26667<br>0.4879 | -0.11667<br>0.7650 | 0.25000<br>0.5165 | 0.01667<br>0.9661 |
| **Body mass change final - capture** | 0.13990<br>0.7196 | -0.07869<br>0.8405 | 0.02623<br>0.9466 | -0.36724<br>0.3309 | -0.37598<br>0.3186 | -0.71698<br>0.0297 | -0.38472<br>0.3066 | -0.66452<br>**0.0509** |
| **Neophobia index initial** | 0.25104<br>0.5147 | 0.36515<br>0.3339 | -0.34233<br>0.3672 | 0.54772<br>0.1269 | -0.70747<br>**0.0330** | -0.54772<br>0.1269 | 0.02282<br>0.9535 | -0.45644<br>0.2168 |
| **Neophobia index final** | 0.41079<br>0.2721 | 0.54772<br>0.1269 | -0.41079<br>0.2721 | 0.54772<br>0.1269 | -0.41079<br>0.2721 | -0.54772<br>0.1269 | 0.13693<br>0.7254 | -0.54772<br>0.1269 |
| **Exploration index initial** | -0.28818<br>0.4521 | -0.32208<br>0.3980 | 0.25427<br>0.5091 | -0.16952<br>0.6628 | 0.16952<br>0.6628 | -0.16952<br>0.6628 | 0.42379<br>0.2557 | 0.01695<br>0.9655 |
| **Exploration index final** | -0.05021<br>0.8979 | -0.11716<br>0.7640 | 0.43515<br>0.2418 | 0.29289<br>0.4444 | -0.30126<br>0.4308 | -0.04184<br>0.9149 | 0.60252<br>0.0860 | -0.11716<br>0.7640 |

Table S3: Candidate model set for exploration of the novel environment in adults.

| Candidate models | LL | k | ΔAICc | Weight |
|---|---|---|---|---|
| Treatment\|Sex + *FKBP5*_HPC | -7.03 | 6 | 0 | 0.86 |
| Treatment\|Sex + baseline_CORT | -9.32 | 6 | 4.58 | 0.09 |
| Treatment\|Sex + negative_feedback | -10.27 | 6 | 6.49 | 0.03 |
| Treatment\|Sex + post-dex_CORT | -11.69 | 6 | 9.33 | 0.01 |
| Treatment\|Sex + stress_response | -11.8 | 6 | 9.53 | 0.01 |
| Treatment\|Sex + post-restraint_CORT | -11.83 | 6 | 9.61 | 0.01 |

Table S4: Candidate model set for neophobia in juveniles.

| Candidate models | LL | k | ΔAICc | Weight |
|---|---|---|---|---|
| Treatment + *FKBP5*_post-dex | -1.11 | 3 | 0 | 0.81 |
| Treatment + baseline_CORT | -3.99 | 3 | 5.77 | 0.05 |
| Treatment + negative_feedback | -4.04 | 3 | 5.86 | 0.05 |
| Treatment + post-restraint_CORT | -4.26 | 3 | 6.29 | 0.04 |
| Treatment + post-dex_CORT | -4.27 | 3 | 6.31 | 0.03 |
| Treatment + stress_response | -4.28 | 3 | 6.34 | 0.03 |

Table S5: Correlation matrix for *FKBP5* relative expression in the hippocampus (HPC), hypothalamus (HYPO), in the blood baseline (base), stress-induced (str) and post-dex (dex) samples at capture (capt) and final series (final) and glucocorticoid receptor (GR) relative expression in the hippocampus and hypothalamus in adults.

| | ***FKBP5*** **HPC** | ***FKBP5*** **HYPO** | ***FKBP5*** **base capt** | ***FKBP5*** **str capt** | ***FKBP5*** **dex capt** | ***FKBP5*** **base final** | ***FKBP5*** **str final** | ***FKBP5*** **dex final** | **GR HPC** | **GR HYPO** |
|---|---|---|---|---|---|---|---|---|---|---|
| ***FKBP5*** **HPC** | 1.00000 | -0.13333<br>0.7324 | 0.16667<br>0.6682 | -0.08333<br>0.8312 | 0.55000<br>0.1250 | 0.25000<br>0.5165 | -0.20000<br>0.6059 | 0.08333<br>0.8312 | 0.65000<br>**0.0501** | 0.15000<br>0.7001 |
| ***FKBP5*** **HYPO** | -0.13333<br>0.7324 | 1.00000 | 0.88333<br>**0.0016** | 0.71667<br>**0.0298** | 0.13333<br>0.7324 | 0.36667<br>0.3317 | -0.13333<br>0.7324 | 0.33333<br>0.3807 | -0.31667<br>0.4064 | 0.65000<br>**0.0501** |
| ***FKBP5*** **base capt** | 0.16667<br>0.6682 | 0.88333<br>**0.0016** | 1.00000 | 0.58333<br>0.0992 | 0.40000<br>0.2861 | 0.76785<br>**0.0157** | -0.16667<br>0.6682 | 0.40000<br>0.2861 | -0.05000<br>0.8984 | 0.68333<br>**0.0424** |
| ***FKBP5*** **str capt** | -0.08333<br>0.8312 | 0.71667<br>**0.0298** | 0.58333<br>0.0992 | 1.00000 | 0.21667<br>0.5755 | 0.58333<br>0.0992 | -0.15000<br>0.7001 | 0.36667<br>0.3317 | -0.38333<br>0.3085 | 0.61667<br>0.0769 |
| ***FKBP5*** **dex capt** | 0.55000<br>0.1250 | 0.13333<br>0.7324 | 0.40000<br>0.2861 | 0.21667<br>0.5755 | 1.00000 | 0.55000<br>0.1250 | -0.38333<br>0.3085 | 0.26667<br>0.4879 | 0.00000<br>1.0000 | 0.18333<br>0.6368 |
| ***FKBP5*** **base final** | 0.25000<br>0.5165 | 0.36667<br>0.3317 | 0.43333<br>0.2440 | 0.58333<br>0.0992 | 0.55000<br>0.1250 | 1.00000 | -0.15000<br>0.7001 | 0.73333<br>**0.0246** | -0.43333<br>0.2440 | 0.05000<br>0.8984 |
| ***FKBP5*** **str final** | -0.20000<br>0.6059 | -0.13333<br>0.7324 | -0.16667<br>0.6682 | -0.15000<br>0.7001 | -0.38333<br>0.3085 | -0.15000<br>0.7001 | 1.00000 | 0.38333<br>0.3085 | -0.25000<br>0.5165 | -0.35000<br>0.3558 |
| ***FKBP5*** **dex final** | 0.08333<br>0.8312 | 0.33333<br>0.3807 | 0.40000<br>0.2861 | 0.36667<br>0.3317 | 0.26667<br>0.4879 | 0.73333<br>**0.0246** | 0.38333<br>0.3085 | 1.00000 | -0.46667<br>0.2054 | -0.25000<br>0.5165 |
| **GR HPC** | 0.65000<br>**0.0501** | -0.31667<br>0.4064 | -0.05000<br>0.8984 | -0.38333<br>0.3085 | 0.00000<br>1.0000 | -0.43333<br>0.2440 | -0.25000<br>0.5165 | -0.46667<br>0.2054 | 1.00000 | 0.25000<br>0.5165 |
| **GR HYPO** | 0.15000<br>0.7001 | 0.65000<br>**0.0501** | 0.68333<br>**0.0424** | 0.61667<br>0.0769 | 0.18333<br>0.6368 | 0.05000<br>0.8984 | -0.35000<br>0.3558 | -0.25000<br>0.5165 | 0.25000<br>0.5165 | 1.00000 |

Table S6: Correlation matrix for *FKBP5* relative expression in the hippocampus (HPC), hypothalamus (HYPO), in the blood baseline (base), stress-induced (str) and post-dex (dex) samples at capture (capt) and final series (final) and glucocorticoid receptor (GR) relative expression in the hippocampus and hypothalamus in juveniles.

| | ***FKBP5*** **HPC** | ***FKBP5*** **HYPO** | ***FKBP5*** **base capt** | ***FKBP5*** **str capt** | ***FKBP5*** **dex capt** | ***FKBP5*** **base final** | ***FKBP5*** **str final** | ***FKBP5*** **dex final** | **GR HPC** | **GR HYPO** |
|---|---|---|---|---|---|---|---|---|---|---|
| ***FKBP5*** **HPC** | 1.00000 | 0.81667<br>**0.0072** | -0.08333<br>0.8312 | 0.36667<br>0.3317 | -0.20000<br>0.6059 | -0.23333<br>0.5457 | -0.06667<br>0.8647 | -0.53333<br>0.1392 | 0.58333<br>0.0992 | 0.63333<br>0.0671 |
| ***FKBP5*** **HYPO** | 0.81667<br>**0.0072** | 1.00000 | -0.41667<br>0.2646 | 0.63333<br>0.0671 | -0.41667<br>0.2646 | -0.01667<br>0.9661 | -0.16667<br>0.6682 | -0.45000<br>0.2242 | 0.33333<br>0.3807 | 0.83333<br>**0.0053** |
| ***FKBP5*** **base capt** | -0.08333<br>0.8312 | -0.41667<br>0.2646 | 1.00000 | -0.28333<br>0.4600 | 0.05000<br>0.8984 | 0.23333<br>0.5457 | 0.33333<br>0.3807 | 0.18333<br>0.6368 | 0.33333<br>0.3807 | -0.30000<br>0.4328 |
| ***FKBP5*** **str capt** | 0.36667<br>0.3317 | 0.63333<br>0.0671 | -0.28333<br>0.4600 | 1.00000 | -0.40000<br>0.2861 | 0.25000<br>0.5165 | 0.45000<br>0.2242 | 0.10000<br>0.7980 | 0.01667<br>0.9661 | 0.66667<br>**0.0499** |
| ***FKBP5*** **dex capt** | -0.20000<br>0.6059 | -0.41667<br>0.2646 | 0.05000<br>0.8984 | -0.40000<br>0.2861 | 1.00000 | 0.21667<br>0.5755 | 0.31667<br>0.4064 | 0.31667<br>0.4064 | -0.20000<br>0.6059 | -0.56667<br>0.1116 |
| ***FKBP5*** **base final** | -0.23333<br>0.5457 | -0.01667<br>0.9661 | 0.23333<br>0.5457 | 0.25000<br>0.5165 | 0.21667<br>0.5755 | 1.00000 | 0.26667<br>0.4879 | 0.70000<br>**0.0358** | -0.28333<br>0.4600 | 0.20000<br>0.6059 |
| ***FKBP5*** **str final** | -0.06667<br>0.8647 | -0.16667<br>0.6682 | 0.33333<br>0.3807 | 0.45000<br>0.2242 | 0.31667<br>0.4064 | 0.26667<br>0.4879 | 1.00000 | 0.35000<br>0.3558 | 0.13333<br>0.7324 | -0.23333<br>0.5457 |
| ***FKBP5*** **dex final** | -0.53333<br>0.1392 | -0.45000<br>0.2242 | 0.18333<br>0.6368 | 0.10000<br>0.7980 | 0.31667<br>0.4064 | 0.70000<br>**0.0358** | 0.35000<br>0.3558 | 1.00000 | -0.55000<br>0.1250 | -0.03333<br>0.9322 |
| **GR HPC** | 0.58333<br>0.0992 | 0.33333<br>0.3807 | 0.33333<br>0.3807 | 0.01667<br>0.9661 | -0.20000<br>0.6059 | -0.28333<br>0.4600 | 0.13333<br>0.7324 | -0.55000<br>0.1250 | 1.00000 | 0.10000<br>0.7980 |
| **GR HYPO** | 0.63333<br>0.0671 | 0.83333<br>**0.0053** | -0.30000<br>0.4328 | 0.66667<br>**0.0499** | -0.56667<br>0.1116 | 0.20000<br>0.6059 | -0.23333<br>0.5457 | -0.03333<br>0.9322 | 0.10000<br>0.7980 | 1.00000 |
